# Extensive genetic diversity among populations of the malaria mosquito *Anopheles moucheti* revealed by population genomics

**DOI:** 10.1101/068247

**Authors:** Caroline Fouet, Colince Kamdem, Stephanie Gamez, Bradley J. White

## Abstract

Malaria vectors are exposed to intense selective pressures due to large-scale intervention programs that are underway in most African countries. One of the current priorities is therefore to clearly assess the adaptive potential of Anopheline populations, which is critical to understand and anticipate the response mosquitoes can elicit against such adaptive challenges. The development of genomic resources that will empower robust examinations of evolutionary changes in all vectors including currently understudied species is an inevitable step toward this goal. Here we constructed double-digest Restriction Associated DNA (ddRAD) libraries and generated 6461 Single Nucleotide Polymorphisms (SNPs) that we used to explore the population structure and demographic history of wild-caught *Anopheles moucheti* from Cameroon. The genome-wide distribution of allelic frequencies among samples best fitted that of an old population at equilibrium, characterized by a weak genetic structure and extensive genetic diversity, presumably due to a large long term effective population size. Estimates of *F*_ST_ and Linkage Disequilibrium (LD) across SNPs reveal a very low genetic differentiation throughout the genome and the absence of segregating LD blocks among populations, suggesting an overall lack of local adaptation. Our study provides the first investigation of the genetic structure and diversity in *An. moucheti* at the genomic scale. We conclude that, despite a weak genetic structure, this species has the potential to challenge current vector control measures and other rapid anthropogenic and environmental changes thanks to its great genetic diversity.

## 1. Introduction

Despite having a widely acknowledged epidemiological significance, most African malaria mosquitoes are so-called “neglected vectors” because the efforts devoted to their study and control are clearly insufficient. *Anopheles moucheti sensu lato* is one of the best examples. This mosquito vector is a group of three related species (*An. moucheti moucheti, An. moucheti nigeriensis*, and *An. moucheti bervoetsi*) distributed across the equatorial forest and distinguishable from each other by slight morphological differences (Kengne et al., 2007). The nominal species of the group, *An. moucheti moucheti* (hereafter *An. moucheti*), is a very efficient and anthropophilic vector especially in rural areas where the highest malaria burden due to *Plasmodium falciparum* infections are recorded (Antonio-Nkondjio et al., 2009, 2008, 2002). In such settings, abundant populations of *An. moucheti* breed year-round in slow moving streams and rivers and often outcompete other main malaria mosquitoes. Despite this epidemiological significance, the evolutionary history and the adaptive potential of this vector remain understudied. Early investigations of the genetic structure based on allozymes and microsatellites showed a significant genetic differentiation among samples from three different countries (Antonio-Nkondjio et al., 2008), but detected little divergence within populations from the same country (Antonio-Nkondjio et al., 2007, 2002). Precisely, very low levels of genetic differentiation were found between populations from Cameroon across eight microsatellite loci, suggesting extensive gene flow at such geographic scales, but detailed studies in other countries are still lacking to fully support this hypothesis. On the other hand, African anopheline populations are increasingly exposed to strong selective pressures associated with insecticide-based malaria control campaigns that have been recently intensified (World Health Organization, 2013). Such pressures represent particularly efficient driving forces that often contribute to the rapid diversification of vector populations in a few decades (Clarkson et al., 2014; Kamdem et al., Unpublished data a; Norris et al., 2015). As a result, a detailed characterization of the genomic architecture of all vectors is important for a critical appraisal of the impacts of malaria control efforts. In this framework, we set out to perform the first genome-wide investigation of natural polymorphism in *An. moucheti*. One of our main goals was to know to what extent assessing the genetic diversity could provide clues about the spatial distribution and help predict the environmental resilience of this species. In principle, evolutionary responses of species to human-induced or natural changes rely largely on available heritable variation, which reflects the evolutionary potential and adaptability to novel environments (Orr and Unckless, 2008). Therefore, the screening of genome-wide variation is supposed to be a sensible approach that may provide a generalized measure of evolutionary potential in species like *An. moucheti* for which direct ecological, evolutionary or functional tests are impossible (Harrisson et al., 2014).

Thanks to recent progresses in sequencing technology, high-resolution sequence information can be generated for virtually any living organism. These technological advances are extraordinary helpful for non-model species with limited genomic resources like mosquitoes (Ellegren, 2014). However, at the exception of *Anopheles gambiae* for which significant genomic studies have been carried out using high-quality sequencing data (Fontaine et al., 2015; Kamdem et al., Unpublished data a; O’Loughlin et al., 2014), the other African malaria vectors have yet to fully benefit from the explosive growth of methods for assessing genetic variation at a fine scale. These neglected vectors face a vicious cycle whereby the lack of basic genomic resources that are critical to generate high-quality sequencing information and to enable robust interpretations of natural polymorphisms greatly contributes to their marginalization. One typical example is *An. moucheti*, which lacks all the vital resources ranging from a laboratory strain, a reference genome assembly, and a physical or linkage map.

To start filling this gap and to shed some light on the evolutionary history and adaptive potential of this vector, we have performed a high-throughput sequencing of reduced representation libraries in 98 wild-caught individuals from Cameroon and identify thousands of RAD loci scattered throughout the genome. Using high-quality Single Nucleotide Polymorphisms (SNPs) identified within these loci, we have investigated the genetic structure of populations and scan genomes of our samples to detect footprints of local adaptation and natural selection. We found that, in our study zone, populations of *An. moucheti* are characterized by a great genetic diversity and extensive gene flow. We argue that this vector is particularly adapted to challenge the selective pressures imposed by vector controls and rapid environmental modifications.

## 2. Material and methods

### 2.1 Mosquito sampling and sequencing

This study included two *An. moucheti* populations from the Cameroonian equatorial forest. A total of 98 mosquitoes (97 adults and 1 larva) were collected in August and November 2013 from Olama and Nyabessan, respectively (Table 1). The two locations are separated by ~200 km (Fig. 1A) and are crossed respectively by the Nyong and the Ntem rivers that provide the breeding sites for *An. moucheti* larvae. Specimens were identified as *An. moucheti moucheti* using morphological identification keys (Gillies and Coetzee, 1987; Gillies and De Meillon, 1968) and a diagnostic PCR, which targets mutations on the ribosomal DNA (Kengne et al., 2007). We extracted genomic DNA using the DNeasy Blood and Tissue kit (Qiagen) for larvae and the Zymo Research MinPrep kit for adult mosquitoes. We used 10µl (~50ng) of genomic DNA to prepare double-digest Restriction-site Associated DNA libraries following a modified protocol of Peterson et al., 2012. *MluC1* and *NlaIII* restriction enzymes were used to digest DNA of individual mosquitoes, yielding RAD-tags of different sizes to which short unique DNA sequences (barcodes and adaptors) were ligated to enable the identification of reads belonging to each specimen. The digestion products were purified and pooled. DNA fragments of around 400bp were selected and amplified via PCR. The distribution of fragment sizes was checked on a BioAnalyzer (Agilent Technologies, Inc., USA) before sequencing. The sequencing was performed on an Illumina HiSeq2000 platform (Illumina Inc., USA) (Genomic Core Facility, University of California, Riverside) to yield single-end reads of 101bp.

**Figure 1:**
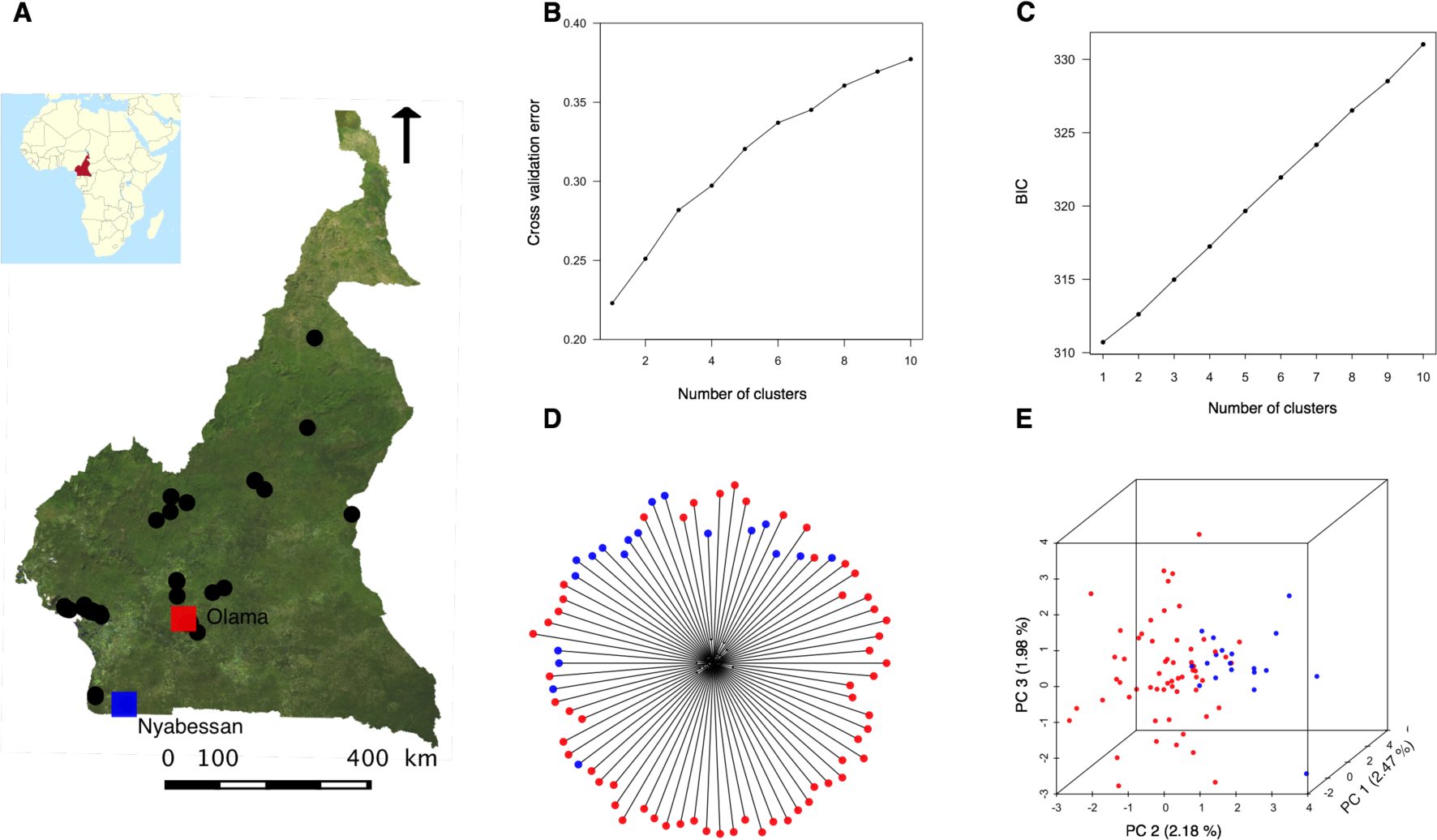
Relationship between *An. moucheti* individuals from Olama and Nyabessan. (A) Map of the study site showing both the locations surveyed (small black dots) and the two villages (large red and blue squares) where *An. moucheti* samples were collected. (B) and (C) Plots of the ADMIXTURE cross-validation error and the Bayesian Information Criterion (BIC) (DAPC) as a function of the number of genetic clusters indicating that k = 1. The lowest BIC and CV error indicate the suggested number of clusters. (D) and (E) Absence of genetic structure within populations illustrated by neighbor-joining and PCA. The percentage of variance explained by each PCA axis is indicated.

**Table 1:** Information on *An. moucheti* samples included in this study.

| Sampling<br>locations | Geographic<br>coordinates | Sampling<br>methods |  |  | Total |
| --- | --- | --- | --- | --- | --- |
|  |  | HLC-OUT | HLC-IN | LC |  |
| Nyabessan | 2°24'00"N, 10°24'00"E | 21 | 15 | 1 | 37 |
| Olama | 3°26'00"N, 11°17'00"E | 30 | 31 | 0 | 61 |
| Total |  |  |  |  | 98 |

HLC-OUT, human landing catches performed outdoor; HLC-IN, human landing catches performed indoor; LC, larval collection

### 2.2 SNP discovery and genotyping

We used the bioinformatics pipeline Stacks v1.35 (Catchen et al., 2013) to process Illumina short reads. The program *process_radtags* was first used to sort the reads according to the barcodes and to trim all reads to 96bp in length by removing index and barcode sequences from the ends of the reads. Reads with ambiguous barcodes, those that did not contain the *NlaIII* recognition site and those with low-quality scores (average Phred score < 33) were excluded. The program *ustacks* was then utilized to perform a *de novo* assembly (i.e., the assembly of reads in “stacks” enabling the creation of consensus RAD loci without prior alignment onto a reference genome sequence) (Catchen et al., 2013, 2011) in each individual in our populations. We allowed a maximum of 2 nucleotide mismatches between stacks (M parameter in *ustacks*) and we required a minimum of three reads to create a stack (m parameter in *ustacks*). Using the *cstacks* program, a catalogue of loci was built to synchronize variations across all individuals in our populations. Finally, we utilized the *populations* program to calculate population genetic parameters and output SNPs in different formats. To avoid bias associated with less informative SNPs or possible false positive SNPs (due to sequencing or pipeline errors), only RAD loci scored in at least 70-75% of individuals were retained for further analyses.

### 2.3 Population genomic analyses

SNP files outputted by the *populations* program were used to assess the population genetic structure with a Principal Component Analysis (PCA) and a Neighbor-Joining (NJ) tree analysis using respectively the R packages *adegenet* and *ape* (Jombart, 2008; Paradis et al., 2004; R Development Core Team, 2008). We also explored patterns of ancestry and admixture among individuals in ADMIXTURE v1.23 (Alexander et al., 2009) with 10-fold cross-validation for k assumed ancestral populations (k= 1 through 6). The optimal number of clusters was confirmed using the Discriminant Analysis of Principal Component (DAPC) method, which explores the number of genetically distinct groups by running a k-means clustering sequentially with increasing numbers and by comparing different clustering solutions using Bayesian Information Criterion (BIC) (Jombart, 2008). We examined the population genetic diversity, conformity to Hardy-Weinberg equilibrium and demographic background using several statistics calculated with the *populations* program. Precisely, to assess the global genetic diversity per population, we calculated the overall nucleotide diversity (π) and the frequency of polymorphic sites within population. To make inferences on the demographic history and to test for departures from Hardy–Weinberg equilibrium, we used the allele frequency spectrum and the Wright’s inbreeding coefficient (*F*_IS_). To quantify the geographic and genetic differentiation between allopatric populations, we estimated the genome-wide average *F*_ST_ (Weir and Cockerham, 1984) on 2000 randomly selected SNPs in Genodive v1.06 (Meirmans and Van Tienderen, 2004). We also conducted an hierarchical Analysis of Molecular Variance (AMOVA) (Excoffier et al., 1992) on the same SNP set to quantify the effects of the geographic origin on the genetic variance among individuals. The statistical significance of *F*_ST_ and AMOVA was assessed with 10000 permutations. Finally, to have a detailed picture of the genomic architecture of divergence, we inspected the genome-wide distribution of locus-specific estimates of *F*_ST_.

### 2.4 Identification of segregating polymorphic chromosomal inversions

In structured *Anopheles* populations whose ecological/genetic divergence is due to polymorphic chromosomal inversions, high values of *F*_ST_ are expected between divergent populations within inversion loci, a pattern consistent with local adaptation of alternative karyotypes (Ayala and Coluzzi, 2005). This is the case for most populations of the main African malaria vectors *An. funestus* and *An. gambiae*, which depict multiple inversion clines in nature (Ayala et al., 2011; Fouet et al., Unpublished data; Kamdem et al., Unpublished data a; O’Loughlin et al., 2014). In addition to scanning genomes of our individuals to identify outlier values of *F*_ST_ that are indicators of selection and local adaptation, we also used Linkage Disequilibrium (LD) analysis to search for the presence of LD blocks corresponding to putative inversion polymorphisms. LD (the nonrandom association of alleles at different loci) provides information about past events and is affected by local adaptation and geographical structure, the demographic history, or the magnitude of selection and recombination across the genome (Lewontin and Kojima, 1960). Notably, high LD is expected in regions bearing inversions relative to the rest of the genome because the neutral recombination rate is notoriously reduced within inversions (Kirkpatrick and Barton, 2006). Thus, assessing genome-wide patterns of LD can reveal clusters of strongly correlated SNPs (LD blocks) corresponding potentially to chromosomal inversions. The R package LDna (Kemppainen et al., 2015) allows the examination of the distinct LD network clusters within the genome of non-model species without the need of a linkage map or reference genome. We have calculated LD, estimated as the *r*^2^ correlation coefficient between all pairs of SNPs, in PLINK v1.09 (Purcell et al., 2007). To avoid spurious LD due to the strong correlation between SNPs located on the same RAD locus, we randomly selected only one SNP within each RAD locus resulting in a dataset of 2569 variants containing less than 15% missing data. LDna was then used to identify LD blocks whose population genetic structure was examined with a PCA.

## 3. Results

### 3.1 *De novo* assembly

In total, 518,218 unique 96-bp RAD loci were identified from *de novo* assembly of reads in 98 individuals. We retained 946 loci that were present in all sampled populations and in at least 75% of individuals in every population, and we identified 3027 high-quality biallelic SNPs from these loci.

### 3.2 Population genetic structure

First, we tested for the presence of cryptic genetic subdivision within *An. moucheti* with PCA, NJ trees and the ADMIXTURE ancestry model. A NJ tree constructed from a matrix of Euclidian distance using allele frequencies at 3027 genome-wide SNPs showed a putative subdivision of *An. moucheti* populations in two genetic clusters (Fig. S1A). The first three axes of PCA also revealed a number of outlier individuals separated from a main cluster (Fig. S1B). However, when we ranked our sequenced individuals based on the number of sequencing reads, we noticed that one of the putative genetic clusters corresponded to a group of individuals having the lowest sequencing coverage (Fig. S1 and Table S1). We excluded all these individuals and reduced our dataset to 78 individuals. We conducted a new *de novo* assembly and analyzed the relationship between the 78 remaining individuals at 6461 SNPs present in at least 70% of individuals using PCA, NJ trees and ADMIXTURE. Both the k-means clustering (DAPC) and the variation of the cross-validation error as a function of the number of ancestral populations in ADMIXTURE revealed that the polymorphism of *An. moucheti* resulted from only one ancestral population (k = 1) (Fig. 1B and 1C). PCA and NJ depicted a homogeneous cluster comprising all 78 individuals providing additional evidence of the lack of genetic or geographic structuring among populations (Fig. 1D and 1E). Unsurprisingly, the overall *F*_ST_ was remarkably low between populations from the two sampling locations Olama and Nyabessan (*F*_ST_ = 0.008, p < 0.005). Similarly, the distribution of *F*_ST_ values across 6461 SNPs showed a large dominance of very low *F*_ST_ values throughout the genome (Fig. 2). The highest per locus *F*_ST_ was only 0.126, while 5006 of the 6461 loci revealed *F*_ST_ near zero. The modest geographic differentiation was also well illustrated by a hierarchical AMOVA, which showed that the genetic variance was explained essentially by within-individual variations (99.7%). Finally, we found very low overall Wright’s inbreeding coefficient (*F*_IS_= 0.0014, p < 0.005 in Nyabessan and *F*_IS_ = 0.0025, p < 0.005 in Olama) (Table 2) suggesting that allelic frequencies within both populations were in accordance with proportions expected under the Hardy-Weinberg equilibrium.

**Figure 2:**
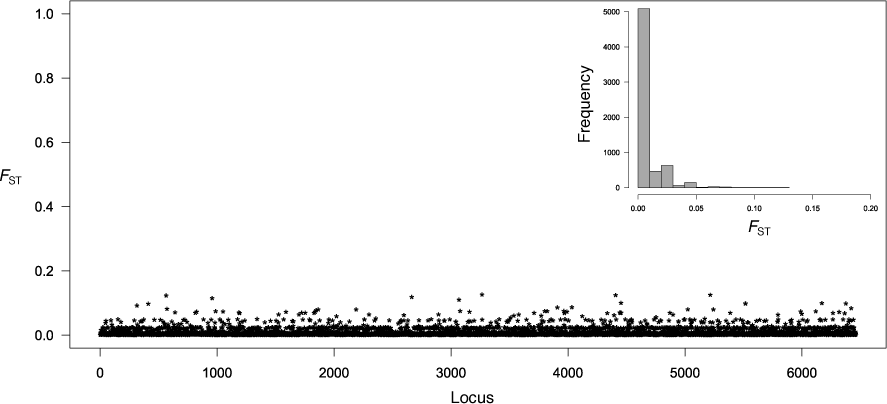
Frequency distribution of *F*_ST_ between Olama and Nyabessan across 6461 SNP loci and plot of these *F*_ST_ values along arbitrary positions in the genome.

**Table 2:**
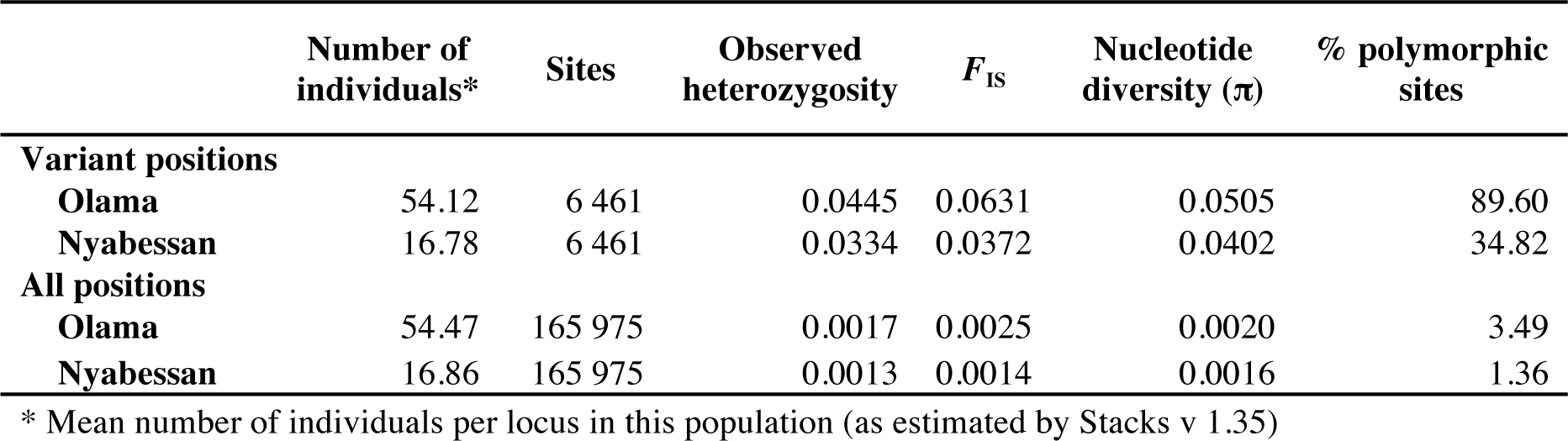
Population genomic parameters based on 6461 variant sites reflecting the genetic diversity and conformity to Hardy-Weingberg equilibrium.

| | Number of<br>individuals* | Sites | Observed<br>heterozygosity | $F_{IS}$ | Nucleotide<br>diversity ($\pi$) | % polymorphic<br>sites |
| --- | --- | --- | --- | --- | --- | --- |
| **Variant positions** |  |  |  |  |  |  |
| **Olama** | 54.12 | 6 461 | 0.0445 | 0.0631 | 0.0505 | 89.60 |
| **Nyabessan** | 16.78 | 6 461 | 0.0334 | 0.0372 | 0.0402 | 34.82 |
| **All positions** |  |  |  |  |  |  |
| **Olama** | 54.47 | 165 975 | 0.0017 | 0.0025 | 0.0020 | 3.49 |
| **Nyabessan** | 16.86 | 165 975 | 0.0013 | 0.0014 | 0.0016 | 1.36 |

\* Mean number of individuals per locus in this population (as estimated by Stacks v 1.35)

### 3.3 Genetic diversity and demographic history

The estimates of the overall nucleotide diversity (π = 0.0020 and π = 0.0016, respectively, in Olama and Nyabessan) (Table 2) were within the range of average values found in other African *Anopheles* species using RADseq approaches (Fouet et al., Unpublished data; Kamdem et al., Unpublished data (a, b); O’Loughlin et al., 2014). Notorious demographic expansions have been described in natural populations of this insect clade (Donnelly et al., 2001), and the values of π observed in *An. moucheti* likely reflect the level of genetic diversity of a population with large effective size. The great genetic diversity of *An. moucheti* was also illustrated by the percentage of polymorphic sites. Of the 6461 variant sites, 89.60% were polymorphic in Olama and 34.82% in Nyabessan (Table 2). The difference observed between the two locations can be related to the sample size (n = 19 in Nyabessan and n = 59 in Olama) or to demographic particularities that persists between the two geographic sites despite a massive gene flow. To infer the demographic history of *An. moucheti*, we examined the Allele Frequency Spectrum (AFS), summarized as the distribution of the major allele in one population. This approach was a surrogate to model-based methods that provide powerful examinations of the history of genetic diversity by modeling the AFS at genome-wide SNP variants, but that couldn’t be implemented here due to the lack of a reference genome assembly. The frequency distribution of the major allele p (Fig. 3) indicates that the majority of polymorphic loci are highly frequent in Olama and Nyabessan as shown by the predominance of SNPs at frequencies equal to 1. Ranges of allele frequencies are also similar in both locations (between 0.47 and 1 in Olama and between 0.34 and 1 in Nyabessan). These frequency ranges are expected for old populations at equilibrium capable of accumulating high amount of genetic diversity.

**Figure 3:**
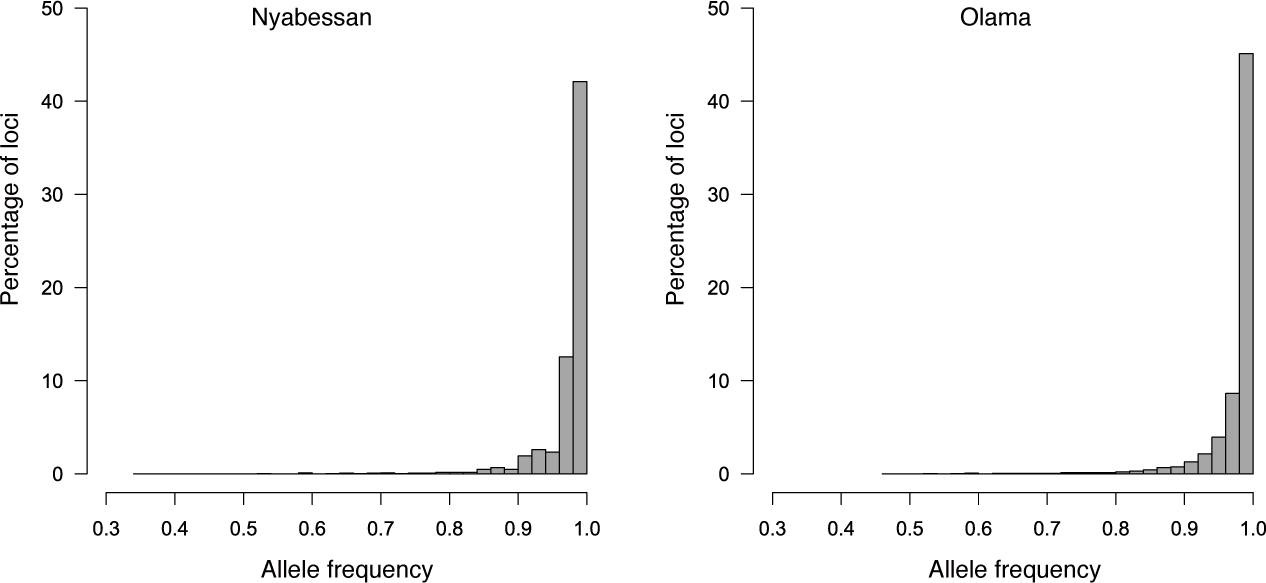
Allele Frequency Spectrum for 6461 SNP loci in Nyabessan and Olama populations. The x-axis presents the frequency of the major allele and the y-axis the frequency distribution of loci in each class of the major allele frequency.

### 3.4 Polymorphic chromosomal inversions and local adaptation

When paracentric inversions are involved in local adaptation, high values of genetic divergence are often observed within inversion loci in natural populations. Cytogenetic analyses of the polytene chromosome of *An. moucheti* have identified three polymorphic chromosomal inversions within samples collected from the sites we have studied (Sharakhova et al., 2014). However, the weak overall population structure and the very low *F*_ST_ values we have detected throughout the genome are clear indicators of the absence of local adaptation. Interestingly, this finding also suggests that none of the polymorphic inversions described previously is actually segregating among our samples, as high values of *F*_ST_ are absent even within inversion loci. We provided further support to this hypothesis by performing LD analyses. First, we found a globally low LD in the *An. moucheti* genome (average genome-wide *r^2^* = 0.0149) as expected in highly polymorphic populations with large effective size. We next used LDna to cluster the LD values and to identify Single Outlier Clusters (SOC) that can be associated with distinct or multiple evolutionary phenomena in the *An. moucheti* history. We set the parameters to collect and screen a high number of SOCs using 2569 highly filtered SNPs, which allowed us to identify 20 independent LD blocks in our samples (Fig. 4). In principle, when these blocks are associated with important events in the evolutionary history of a species, downstream analyses can reveal clear pattern reflecting the underlying process (Kemppainen et al., 2015). This has been illustrated for example by studies demonstrating that SNPs within SOCs generated by polymorphic inversions in *Anopheles baimaii* clearly separate the three expected karyotypes (inverted homozygotes, heterozygotes and standard homozygotes) (Kemppainen et al., 2015). We conducted downstream analyses with a PCA using SNPs identified within the SOCs. As shown in Fig S2, although individuals were occasionally spread along three PCA axes, no distinct cluster could be identified from any of the 20 SOCs. These results were consistent with the absence of segregating inversions and local adaptation in our samples and corroborated low *F*_ST_ values observed throughout the genome. Precisely, in our data, we couldn’t identify polymorphic inversions whose karyotype frequencies change between Olama and Nyabessan due to a differential adaption between the two sites. Some of the different SOCs identified can be associated with other processes that were not captured by our analytical approach; others are probably methodological artifacts associated with the LDna pipeline (Kemppainen et al., 2015).

**Figure 4:**
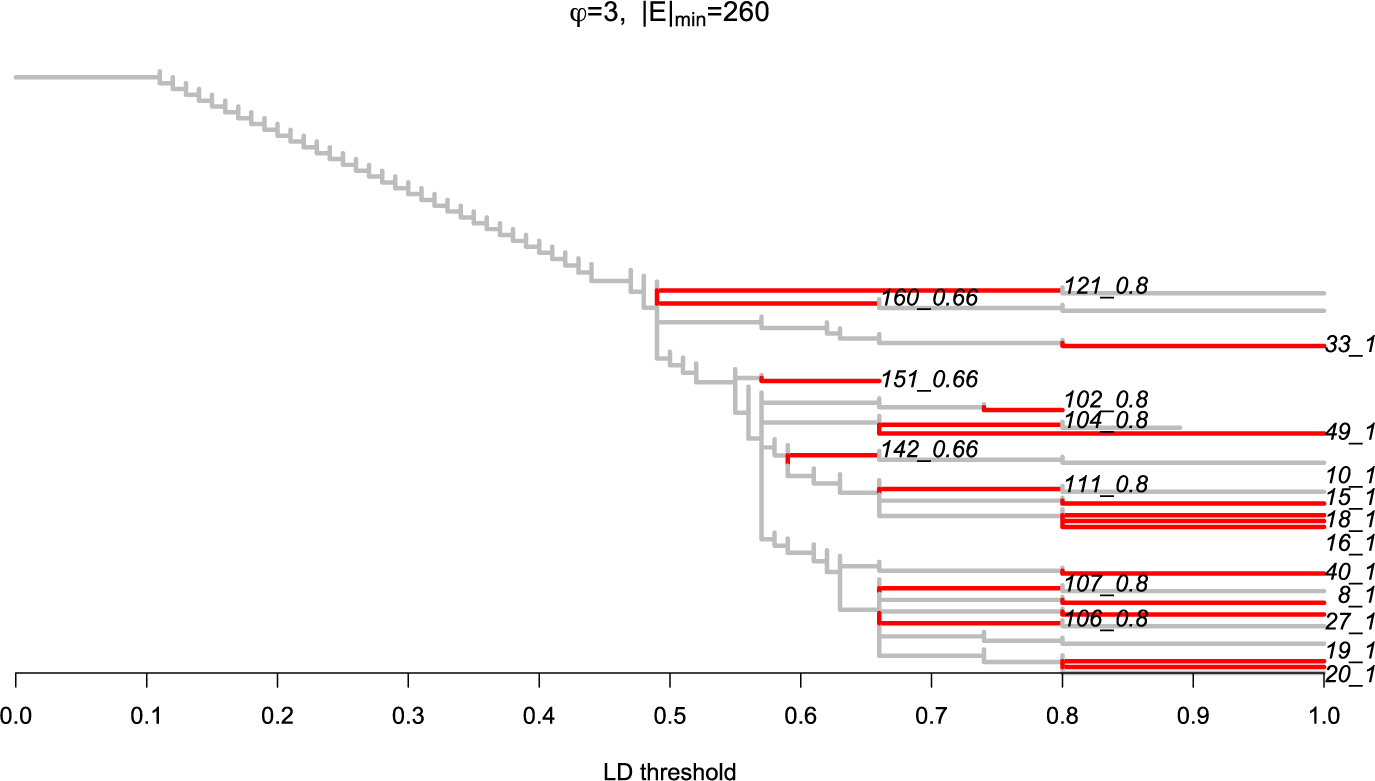
LDna analyses on 2569 SNPs showing the presence of 20 Single Outlier Clusters (SOCs) of linkage disequilibrium in *An. moucheti*. The graph presents the results obtained with values of the two parameters: ϕ (which controls when clusters are defined as outliers) and |*E*|_min_, the minimum number of edges required for a LD cluster to be considered as an outlier, indicated on top. LD thresholds are shown on the x-axis.

## 4. Discussion

We have analyzed the genome-wide polymorphism and characterized some of the baseline population genomic parameters in *An. moucheti*, an important malaria vector in rural areas across the African rainforest. We found very little differentiation among our samples, with most of the genetic variation distributed within individuals. Although a more substantial sampling will be necessary to fully dissect the population genetic structure of this species, our finding likely reflects the current dynamic of *An. moucheti* populations in Cameroon. It is worth mentioning that we have surveyed a total of 28 locations across the country (Fig 1A), some of which were known from several past surveys to harbor *An. moucheti* populations (Antonio-Nkondjio et al., 2013, 2009, 2008, 2006, 2002; Kengne et al., 2007), but we confirmed the presence of the species in only 2 villages. Extant populations of *An. moucheti* are distributed in patches of favorable habitats along river networks where larval populations breed. Our results indicate that despite this apparent fragmentation, connectivity and gene flow are high among population aggregates. The weak population genetic structure of *An. moucheti* observed with genome-wide markers corroborated results obtained with microsatellites and allozymes (Antonio-Nkondjio et al., 2008, 2002). A survey of eight microsatellite loci revealed that the highest *F*_ST_ among Cameroonian populations was as low as 0.003. Nevertheless, a substantial differentiation was found between samples from different countries consistent with an isolation-by-distance model (Antonio-Nkondjio et al., 2008). It is clear that a deep sequencing of continental populations is necessary to further clarify the status of these putative subpopulations. However, samples collected at lower spatial scales like ours are also very relevant as they can allow robust inferences about ongoing selective processes that cannot be captured at continental scale. Although RADseq samples only a small fraction of the genome and certain signatures of selection are likely missing when reduce representation sequencing approaches are used, it has been shown that such approaches can effectively capture strong footprints of selection across genomes of *Anopheles* mosquitoes (Fouet et al., Unpublished data; Kamdem et al., Unpublished data a). We have found that signatures of selection are rare in the genome of *An. moucheti* populations from the Cameroonian rainforest. Populations remain largely undifferentiated throughout the genome, with *F*_ST_ values near zero across the vast majority of variations suggesting that no local adaptation is ongoing. This perception is further supported by the absence of segregating linkage disequilibrium blocks between geographic locations. The characterization of chromosomal inversions with cytogenetic methods can be laborious and challenging (Kirkpatrick, 2010; Sharakhova et al., 2014). So far, three paracentric polymorphic inversions have been discovered in *An. moucheti* in Cameroon (Sharakhova et al., 2014). The ecological, behavioral or functional roles of these inversion polymorphisms remain unknown. We have implemented a recently designed method that uses Next Generation Sequencing and LD estimates to indirectly identify paracentric inversions whose karyotype frequencies varies among populations due to local adaptation (Kemppainen et al., 2015). Our LD analyses revealed the presence of a few LD clusters that are however not associated with inversions. On the other hand, the low overall LD observed across the genome reflected the significant genetic polymorphism that seems to prevail within *An. moucheti* populations. This polymorphism translates into exceptional levels of overall genetic diversity and very high percentage of polymorphic sites that are in the range of values observed in other mosquito species undergoing significant demographic expansions (Donnelly et al., 2001; Fouet et al., Unpublished data; Kamdem et al., Unpublished data a). The amount of neutral genetic diversity is often viewed as a correlate of the adaptive potential of a species (Orr and Unckless, 2008). Although the relationship is more complex in reality, estimates of neutral genetic diversity are commonly used in conservation biology as an intuitive conceptual and management framework to assess the genetic resilience of endangered species (Bonin et al., 2007; Latta IV et al., 2010). Our population genomic analyses have depicted *An. moucheti* as a species with a great genetic diversity and hence a sustainable long-term adaptive resilience. Implications of our findings in malaria epidemiology and control can be very significant. First, *An. moucheti* is essentially endophilic and is particularly sensitive to the principal measures currently employed to control malaria in Sub-Saharan Africa such as the massive use of Insecticide Treated Nets (ITNs) and Indoor Residual insecticide Spraying (IRS). For example, estimates of population effective size in one village in Equatorial Guinea indicated that both mass distribution of ITNs and IRS campaigns resulted in a decline of approximately 55% of *An. moucheti* (Athrey et al., 2012). However, the great genetic diversity and the massive gene flow we observed within populations could easily enable this vector to challenge population declines and recover from shallow bottlenecks. Moreover, most insecticide resistance mechanisms found in insects exploit standing genetic variation to rapidly respond to the evolutionary challenge by increasing the frequency of existing variations rather than relying on infrequent *de novo* mutations (Messer and Petrov, 2013). As a result, despite the current sensitivity of *An. moucheti* to common insecticides, the significant amount of standing genetic variation provides the species with a great potential to challenge insecticide-based interventions and other types of human-induced stress.

## 5. Conclusions

Recent advances in sequencing allow sensitive genomic data to be generated for virtually any species (Ellegren, 2014). However, the most important information we can obtain from population resequencing approaches often depends on the availability and the quality of genomic resources such as a well-annotated reference genome. The reduced genome sequencing strategy (RADseq) offers a cost-effective strategy that can be used to effectively study the genetic variation in a broad range of species from yeast to plants, insects, etc., in the absence of a reference genome. We have extended this approach to the study of the genetic structure of an understudied mosquito species with a great epidemiological significance. We have provided both significant baseline population genomic data and the methodological validation of one approach that should motivate further studies on this species and other understudied anopheline mosquitoes lacking genomic resources.

## Acknowledgements

Funding for this project was provided by the University of California Riverside and NIH grants 1R01AI113248 and 1R21AI115271 to BJW. We thank populations and authorities of the locations surveyed for their kind collaboration. We thank the anonymous reviewers for their careful reading of our manuscript and their many insightful comments and suggestions.

## Author contributions

Conceived and designed the experiments: CF CK BJW. Performed the experiments: CF CK SG BJW. Analyzed the data: CF CK BJW. Wrote the paper: CF CK BJW.

## Supplemental Material

**Figure S1:**
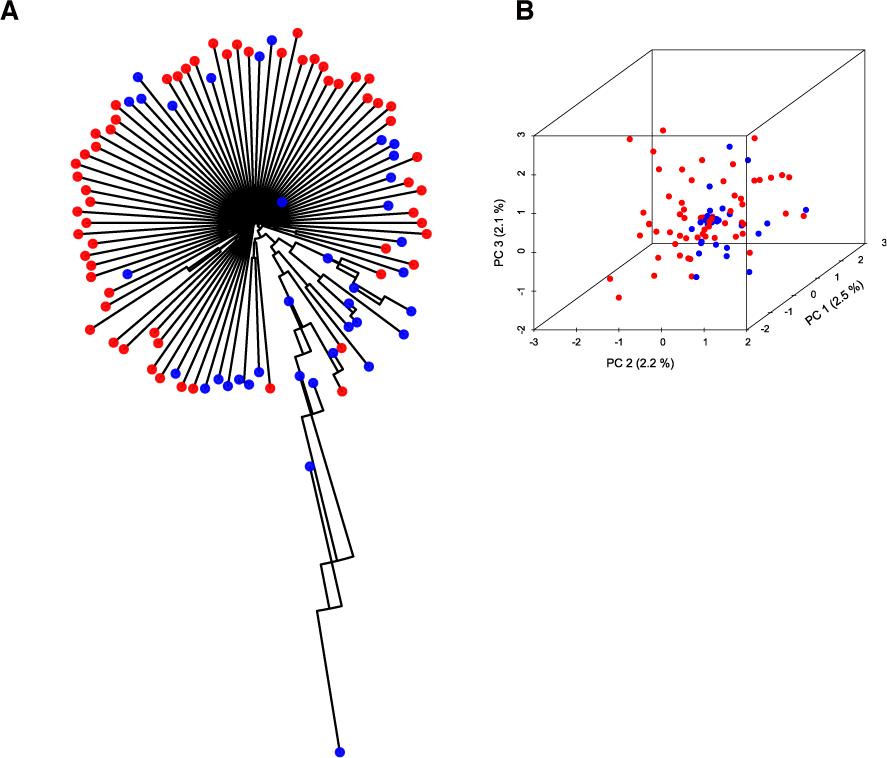
Selection of individuals included in final analyses based on the average per individual sequencing coverage. Neighbor-joining tree (A) and PCA (B) indicating spurious population structure due to individuals with low sequencing coverage in Olama (red) and Nyabessan (blue).

**Figure S2:**
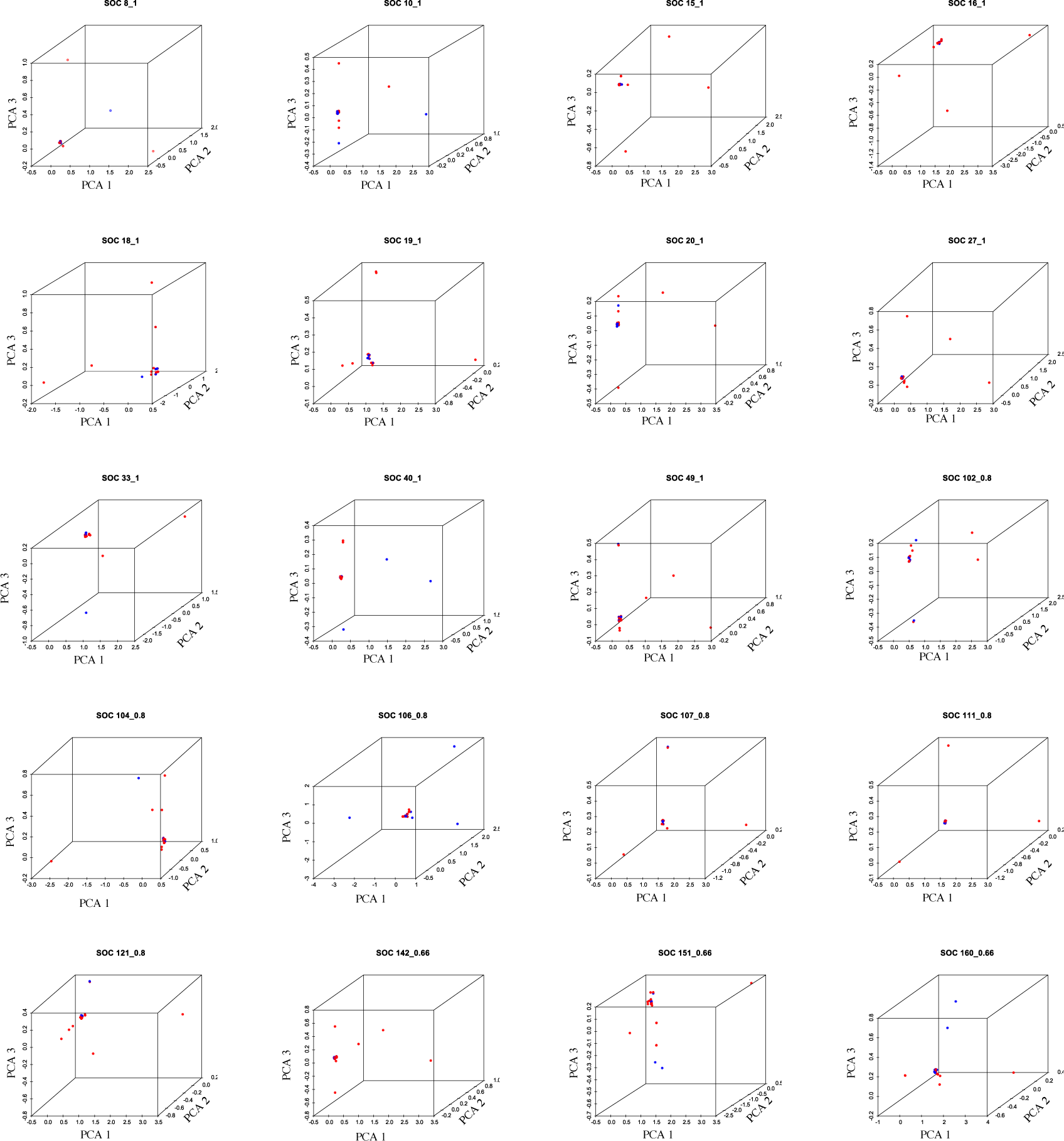
PCA indicating the population genetic structure inferred from SNPs within the 20 Single Outlier Clusters (SOCs) of linkage disequilibrium identified in *An. moucheti* (red: Olama; blue: Nyabessan).

**Table S1:** Distribution of the number of reads among sequenced individuals. Individuals below the dashed line were excluded from analysis.

| Mosquito ID | Total number of reads | Ambiguous/low quality reads | Retained reads | Sampling location |
| --- | --- | --- | --- | --- |
| 43 | 5190686 | 140706 | 5044739 | Olama |
| 850 | 4211705 | 136872 | 4066857 | Nyabessan |
| 27 | 2888406 | 77609 | 2800516 | Olama |
| 97 | 2773623 | 56956 | 2714480 | Olama |
| 94 | 2729590 | 61848 | 2660187 | Olama |
| 26 | 1915405 | 45856 | 1866121 | Olama |
| 82 | 1910651 | 50411 | 1854108 | Olama |
| 37 | 1764370 | 51133 | 1706506 | Olama |
| 95 | 1720323 | 37181 | 1678873 | Olama |
| 93 | 1450516 | 31318 | 1406481 | Olama |
| 25 | 1334643 | 45439 | 1285047 | Olama |
| 79 | 1263586 | 35615 | 1222166 | Olama |
| 80 | 1259353 | 42228 | 1202777 | Olama |
| 98 | 957698 | 20322 | 934945 | Olama |
| 49 | 874718 | 20033 | 852633 | Olama |
| 36 | 867344 | 21542 | 842175 | Olama |
| 99 | 853444 | 17206 | 834357 | Olama |
| 84 | 836989 | 18434 | 816949 | Olama |
| 76 | 850380 | 26481 | 814198 | Olama |
| 31 | 782599 | 18231 | 759965 | Olama |
| 86 | 761099 | 17235 | 741314 | Olama |
| 81 | 735291 | 13915 | 716347 | Olama |
| 1007 | 732960 | 19269 | 710564 | Nyabessan |
| 48 | 725668 | 17270 | 705049 | Olama |
| 743 | 703141 | 15098 | 686817 | Nyabessan |
| 46 | 639840 | 16191 | 621897 | Olama |
| 42 | 634781 | 16180 | 617726 | Olama |
| 92 | 593819 | 12260 | 579932 | Olama |
| 34 | 575175 | 11846 | 561789 | Olama |
| 90 | 579585 | 17480 | 556841 | Olama |
| 868 | 572514 | 16379 | 553831 | Nyabessan |
| 47 | 569810 | 12524 | 551627 | Olama |
| 28 | 557485 | 12363 | 534835 | Olama |
| 30 | 552365 | 13813 | 534263 | Olama |
| 38 | 541076 | 13190 | 525801 | Olama |
| L_627 | 531803 | 12874 | 516440 | Nyabessan |
| 32 | 520421 | 11244 | 507425 | Olama |
| 78 | 510457 | 11112 | 493998 | Olama |
| 19 | 511063 | 15054 | 493676 | Olama |
| 87 | 507445 | 10896 | 492038 | Olama |
| 85 | 498180 | 10747 | 485508 | Olama |
| 792 | 478211 | 10753 | 465355 | Nyabessan |
| 35 | 467060 | 9616 | 455391 | Olama |
| 33 | 463964 | 10001 | 453130 | Olama |
| 60 | 465276 | 11304 | 452748 | Olama |
| 77 | 428787 | 10568 | 415155 | Olama |
| 45 | 391371 | 8565 | 380806 | Olama |
| 61 | 372074 | 10772 | 360550 | Olama |
| 851 | 363537 | 10079 | 352761 | Nyabessan |
| 40 | 346713 | 6421 | 339007 | Olama |
| 89 | 350001 | 12049 | 331665 | Olama |
| 15 | 341218 | 8157 | 331365 | Olama |
| 91 | 337294 | 12007 | 320481 | Olama |
| 44 | 323246 | 7406 | 313719 | Olama |
| 731 | 315382 | 10965 | 302496 | Nyabessan |
| 100 | 311056 | 5531 | 301968 | Olama |
| 729 | 295394 | 6781 | 287687 | Nyabessan |
| 18 | 290682 | 8206 | 281904 | Olama |
| 39 | 225981 | 6085 | 217620 | Olama |
| 73 | 220793 | 6020 | 212919 | Olama |
| 70 | 189567 | 4906 | 183785 | Olama |
| 756 | 188521 | 4639 | 182974 | Nyabessan |
| 75 | 181946 | 6799 | 173364 | Olama |
| 730 | 178968 | 5081 | 171828 | Nyabessan |
| 758 | 177709 | 5115 | 170782 | Nyabessan |
| 732 | 161697 | 4019 | 149839 | Nyabessan |
| 742 | 149159 | 3223 | 144972 | Nyabessan |
| 793 | 132552 | 4456 | 126135 | Nyabessan |
| 14 | 109314 | 3398 | 104900 | Olama |
| 96 | 99548 | 2293 | 95813 | Olama |
| 741 | 94635 | 2614 | 90106 | Nyabessan |
| 724 | 88884 | 3176 | 83605 | Nyabessan |
| 16 | 85486 | 2665 | 81748 | Olama |
| 88 | 83193 | 3414 | 78289 | Olama |
| 777 | 80457 | 2346 | 77695 | Nyabessan |
| 41 | 80127 | 2363 | 75238 | Olama |
| 72 | 76135 | 2366 | 73512 | Olama |
| 723 | 71212 | 2266 | 67258 | Nyabessan |
| 17 | 67037 | 2188 | 64443 | Olama |
| 795 | 65448 | 1778 | 62623 | Nyabessan |
| 869 | 57496 | 2654 | 54157 | Nyabessan |
| 760 | 45620 | 1610 | 43127 | Nyabessan |
| 727 | 43892 | 1402 | 40694 | Nyabessan |
| 71 | 37279 | 1233 | 35811 | Olama |
| 794 | 36669 | 1359 | 34439 | Nyabessan |
| 754 | 35476 | 1137 | 33712 | Nyabessan |
| 726 | 35274 | 1149 | 33230 | Nyabessan |
| 757 | 37431 | 1289 | 32648 | Nyabessan |
| 761 | 29783 | 947 | 28368 | Nyabessan |
| 833 | 27881 | 800 | 26582 | Nyabessan |
| 776 | 24245 | 783 | 22874 | Nyabessan |
| 755 | 24690 | 1012 | 22274 | Nyabessan |
| 753 | 19802 | 1301 | 17379 | Nyabessan |
| 728 | 17829 | 713 | 16750 | Nyabessan |
| 725 | 16069 | 706 | 14676 | Nyabessan |
| 810 | 14088 | 610 | 12963 | Nyabessan |
| 762 | 12307 | 568 | 11575 | Nyabessan |
| 763 | 11562 | 901 | 10374 | Nyabessan |

## References

Alexander, D.H., Novembre, J., Lange, K., 2009. Fast model-based estimation of ancestry in unrelated individuals. Genome Res. 19, 1655–1664.

Antonio-Nkondjio, C., Demanou, M., Etang, J., Bouchite, B., 2013. Impact of cyfluthrin (Solfac EW050) impregnated bed nets on malaria transmission in the city of Mbandjock: lessons for the nationwide distribution of long-lasting insecticidal nets (LLINs) in Cameroon. Parasit. Vectors 6, 10. doi:10.1186/1756-3305-6-10

Antonio-Nkondjio, C., Kerah, C.H., Simard, F., Awono-Ambene, P., Chouaibou, M., Tchuinkam, T., Fontenille, D., 2006. Complexity of the malaria vectorial system in Cameroon: contribution of secondary vectors to malaria transmission. J. Med. Entomol. 43, 1215–1221. doi:10.1603/0022-2585(2006)43[1215:COTMVS]2.0.CO;2

Antonio-Nkondjio, C., Ndo, C., Awono-Ambene, P., Ngassam, P., Fontenille, D., Simard, F., 2007. Population genetic structure of the malaria vector Anopheles moucheti in south Cameroon forest region. Acta Trop. 101, 61–68. doi:10.1016/j.actatropica.2006.12.004

Antonio-Nkondjio, C., Ndo, C., Costantini, C., Awono-Ambene, P., Fontenille, D., Simard, F., 2009. Distribution and larval habitat characterization of Anopheles moucheti, Anopheles nili, and other malaria vectors in river networks of southern Cameroon. Acta Trop. 112, 270–276. doi:10.1016/j.actatropica.2009.08.009

Antonio-Nkondjio, C., Ndo, C., Kengne, P., Mukwaya, L., Awono-Ambene, P., Fontenille, D., Simard, F., 2008. Population structure of the malaria vector Anopheles moucheti in the equatorial forest region of Africa. Malar. J. 7, 120. doi:10.1186/1475-2875-7-120

Antonio-Nkondjio, C., Simard, F., Cohuet, A., Fontenille, D., 2002. Morphological variability in the malaria vector, Anopheles moucheti, is not indicative of speciation: evidences from sympatric south Cameroon populations. Infect. Genet. Evol. 2, 69–72. doi:10.1016/S1567-1348(02)00084-9

Athrey, G., Hodges, T.K., Reddy, M.R., Overgaard, H.J., Matias, A., Ridl, F.C., Kleinschmidt, I., Caccone, A., Slotman, M.a, 2012. The effective population size of malaria mosquitoes: large impact of vector control. PLoS Genet. 8, e1003097. doi:10.1371/journal.pgen.1003097

Ayala, D., Fontaine, M.C., Cohuet, A., Fontenille, D., Vitalis, R., Simard, F., 2011. Chromosomal inversions, natural selection and adaptation in the malaria vector Anopheles funestus. Mol. Biol. Evol. 28, 745–758. doi:10.1093/molbev/msq248

Ayala, F.J., Coluzzi, M., 2005. Chromosome speciation: humans, Drosophila, and mosquitoes. Proc. Natl. Acad. Sci. U. S. A. 102 Suppl, 6535–42. doi:10.1073/pnas.0501847102

Bonin, A., Nicole, F., Pompanon, F., Miaud, C., Taberlet, P., 2007. Population adaptive index: A new method to help measure intraspecific genetic diversity and prioritize populations for conservation. Conserv. Biol. 21, 697–708. doi:10.1111/j.1523-1739.2007.00685.x

Catchen, J., Hohenlohe, P. a., Bassham, S., Amores, A., Cresko, W.., 2013. Stacks: An analysis tool set for population genomics. Mol. Ecol. 22, 3124–3140. doi:10.1111/mec.12354

Catchen, J.M., Amores, A., Hohenlohe, P., Cresko, W., Postlethwait, J.H., 2011. Stacks: building and genotyping Loci de novo from short-read sequences. G3 (Bethesda). 1, 171–82. doi:10.1534/g3.111.000240

Clarkson, C.S., Weetman, D., Essandoh, J., Yawson, A.E., Maslen, G., Manske, M., Field, S.G., Webster, M., Antão, T., MacInnis, B., Kwiatkowski, D., Donnelly, M.J., 2014. Adaptive introgression between Anopheles sibling species eliminates a major genomic island but not reproductive isolation. Nat. Commun. 5, 4248. doi:10.1038/ncomms5248

Donnelly, M.J., Licht, M.C., Lehmann, T., 2001. Evidence for recent population expansion in the evolutionary history of the malaria vectors Anopheles arabiensis and Anopheles gambiae. Mol. Biol. Evol. 18, 1353–1364. doi:10.1093/oxfordjournals.molbev.a003919

Ellegren, H., 2014. Genome sequencing and population genomics in non-model organisms. Trends Ecol. Evol. 29, 51–63. doi:10.1016/j.tree.2013.09.008

Excoffier, L., Smouse, P.E., Quattro, J.M., 1992. Analysis of molecular variance inferred from metric distances among DNA haplotypes: Application to human mitochondrial DNA restriction data. Genetics 131, 479–491. doi:10.1007/s00424-009-0730-7

Fontaine, M.C., Pease, J.B., Steele, a., Waterhouse, R.M., Neafsey, D.E., Sharakhov, I. V., Jiang, X., Hall, a.B., Catteruccia, F., Kakani, E., Mitchell, S.N., Wu, Y.-C., Smith, H. a., Love, R.R., Lawniczak, M.K., Slotman, M.a., Emrich, S.J., Hahn, M.W., Besansky, N.J., 2015. Extensive introgression in a malaria vector species complex revealed by phylogenomics. Science (80). 347, 1258524–1258524. doi:10.1126/science.1258524

Fouet, C., Kamdem, C., White, B.J., 2016. Chromosomal inversions facilitate chromosome-scale evolution in Anopheles funestus. bioRxiv.

Gillies, M.T., Coetzee, M., 1987. A supplement to the Anophelinae of Africa south of the Sahara. The South African Institute for Medical Research, Johannesburg.

Gillies, M.T., De Meillon, B., 1968. The Anophelinae of Africa South of the Sahara, Second Edi. ed. Publications of the South African Institute for Medical Research, Johannesburg.

Harrisson, K.a., Pavlova, A., Telonis-Scott, M., Sunnucks, P., 2014. Using genomics to characterize evolutionary potential for conservation of wild populations. Evol. Appl. n/a-n/a. doi:10.1111/eva.12149

Jombart, T., 2008. adegenet: a R package for the multivariate analysis of genetic markers. Bioinformatics 24, 1403–1405.

Kamdem, C., Fouet, C., Gamez, S., White, B.J., 2016a. Pollutants and insecticides drive local adaptation in African malaria mosquitoes. bioRxiv.

Kamdem, C., Fouet, C., Gamez, S., White, B.J., 2016b. Genomic signatures of introgression at late stages of speciation in a malaria mosquito. bioRxiv.

Kemppainen, P., Knight, C.G., Sarma, D.K., Hlaing, T., Prakash, A., Maung Maung, Y.N., Somboon, P., Mahanta, J., Walton, C., 2015. Linkage disequilibrium network analysis (LDna) gives a global view of chromosomal inversions, local adaptation and geographic structure. Mol. Ecol. Resour. n/a-n/a. doi:10.1111/1755-0998.12369

Kengne, P., Antonio-Nkondjio, C., Awono-Ambene, H.P., Simard, F., Awolola, T.S., Fontenille, D., 2007. Molecular differentiation of three closely related members of the mosquito species complex, Anopheles moucheti, by mitochondrial and ribosomal DNA polymorphism. Med. Vet. Entomol. 21, 177–182. doi:10.1111/j.1365-2915.2007.00681.x

Kirkpatrick, M., 2010. How and why chromosome inversions evolve. PLoS Biol. 8. doi:10.1371/journal.pbio.1000501

Kirkpatrick, M., Barton, N., 2006. Chromosome inversions, local adaptation and speciation. Genetics 173, 419–434. doi:10.1534/genetics.105.047985

Latta IV, L.C., Fisk, D.L., Knapp, R.A., Pfrender, M.E., 2010. Genetic resilience of Daphnia populations following experimental removal of introduced fish. Conserv. Genet. 11, 1737–1745. doi:10.1007/s10592-010-0067-y

Lewontin, R.C., Kojima, K., 1960. The evolutionary dynamics of complex polymorphisms. Evolution (N. Y). doi:10.2307/2405995

Meirmans, P., Van Tienderen, P., 2004. GENOTYPE and GENODIVE: two programs for the analysis of genetic diversity of asexual organisms. Mol. Ecol. Notes 4, 792–794.

Messer, P.W., Petrov, D., 2013. Population genomics of rapid adaptation by soft selective sweeps. Trends Ecol. Evol. 28, 659–669. doi:10.1016/j.tree.2013.08.003

Norris, L.C., Main, B.J., Lee, Y., Collier, T.C., Fofana, A., Cornel, A.J., Lanzaro, G.C., 2015. Adaptive introgression in an African malaria mosquito coincident with the increased usage of insecticide-treated bed nets. Proc. Natl. Acad. Sci. 201418892. doi:10.1073/pnas.1418892112

O’Loughlin, S.M., Magesa, S., Mbogo, C., Mosha, F., Midega, J., Lomas, S., Burt, A., 2014. Genomic Analyses of Three Malaria Vectors Reveals Extensive Shared Polymorphism but Contrasting Population Histories. Mol. Biol. Evol. 1–14. doi:10.1093/molbev/msu040

Orr, H.A., Unckless, R.L., 2008. Population extinction and the genetics of adaptation. Am. Nat. 172, 160–9. doi:10.1086/589460

Paradis, E., Claude, J., Strimmer, K., 2004. Analyses of Phylogenetics and Evolution in R language. Bioinformatics 20, 289–290.

Peterson, B.K., Weber, J.N., Kay, E.H., Fisher, H.S., Hoekstra, H.E., 2012. Double Digest RADseq: An Inexpensive Method for De Novo SNP Discovery and Genotyping in Model and Non-Model Species. PLoS One 7, e37135. doi:10.1371/journal.pone.0037135

Purcell, S., Neale, B., Todd-Brown, K., Thomas, L., Ferreira, M., Bender, D., Maller, J., Sklar, P., de Bakker, P., Daly, M., Sham, P., 2007. PLINK: a toolset for whole-genome association and population-based linkage analysis. Am. J. Hum. Genet. 81.

Sharakhova, M. V., Antonio-Nkondjio, C., Xia, a., Ndo, C., Awono-Ambene, P., Simard, F., Sharakhov, I. V., 2014. Polymorphic chromosomal inversions in Anopheles moucheti, a major malaria vector in Central Africa. Med. Vet. Entomol. 28, 337–340. doi:10.1111/mve.12037

Team, R.D.C., 2008. R: A language and environment for statistical computing. R Foundation for Statistical Computing, Vienna, Austria.

Weir, B.S., Cockerham, C.C., 1984. Estimating F-statistics for the analysis of population structure. Evolution (N. Y). 38, 1358–1370.

World Health Organization, 2013. World malaria report 2013. World Health WHO/HTM/GM, 238. doi:ISBN 978 92 4 1564403

